# Retinal origin of orientation but not direction selective maps in the superior colliculus

**DOI:** 10.1101/2023.08.18.553874

**Authors:** Daniel de Malmazet, Norma K. Kühn, Chen Li, Karl Farrow

## Abstract

Neurons in the mouse superior colliculus (“colliculus”) are arranged in ordered spatial maps. While orientation-selective (OS) neurons form a concentric map aligned to the center of vision, direction-selective (DS) neurons are arranged in patches with changing preferences across the visual field. It remains unclear if these maps are a consequence of feed-forward input from the retina or local computations in the colliculus. To determine whether these maps originate in the retina, we mapped the local and global distribution of orientation- and direction-selective retinal ganglion cell boutons using in-vivo two-photon calcium imaging. We found that OS boutons formed patches that matched the distribution of OS neurons within the colliculus. DS boutons displayed less regional specializations, better reflecting the organization of DS neurons in the retina. Both eyes convey similar orientation but different DS inputs to the colliculus, as shown in recordings from retinal explants. These data demonstrate that orientation and direction maps within the colliculus are independent, implying that orientation maps are inherited from the retina, but direction maps require local circuits.

## Introduction

Visually guided behaviors require coping with distinct sensorymotor relationships in different locations of the visual field. Unsurprisingly, visual field specializations have been observed throughout the visual system of vertebrates. This includes in the retina where photoreceptors and ganglion cells can be asymmetrically spread across the visual field to match an animal’s sensory environment, or behavioral needs ^1–3^. These asymmetries are repeated in higher visual areas including the visual cortex, visual thalamus and superior colliculus (“colliculus”) ^4–7^.

The colliculus is a midbrain region that receives direct input from the retina and uses this information to trigger eye, head and body movements towards or away from objects of interest ^8,9^. Like much of the early visual system the colliculus is retinotopically organized and has prominent regional specializations ^4,10–13^. In rodents, these regional specializations include a clear distinction between the behaviors associated with the processing of visual information in the upper and lower visual fields and the clustered organization of orientation-selective (OS) and direction-selective (DS) neurons ^7,14,15^.

In mice, OS and DS neurons are distributed across the colliculus in a set of patches, within which neurons share similar tuning properties, where the global arrangement of each population is distinct ^10–13,16^. While, OS neurons are arranged in concentric circles around the center of vision ^10^, DS neurons exhibit a sharp transition across the binocular-monocular border, with a broad tendency to prefer upwards-nasal directions in the upper visual field ^11,13^. Broadly, the topographic maps of both, OS and DS neurons, show a correspondence to the visual flow-fields created by self-motion, while DS neurons appear well suited to detect prey in the frontal visual fields, and incoming predators in the upper visual field ^14,17^. This contrasts with the retina, where OS and DS ganglion cells have a predominantly unbiased distribution, where each orientation and direction are equally represented across the visual field ^18,19^. The DS responses of collicular neurons requires input from DS ganglion cells ^20^, and recent modelling work suggests that orientation maps might be formed by spontaneous waves of activity from non-OS ganglion cells during development ^21^. This raises the question at which processing stage the maps of orientation and direction of the colliculus are formed.

In this study we investigate whether the orientation and direction maps in the colliculus originate in the spatial distributions of OS and DS inputs from the retina. To do this, we used two-photon calcium imaging of retinal boutons within the colliculus. We found that while the distribution of OS boutons in the superficial layers matched the spatial distribution of neurons within the colliculus, the properties of DS retinal boutons reflected the spatial distribution observed in the retina. These results suggest that OS retinal ganglion cells (“ganglion cells”) provide regionspecific inputs to the colliculus, playing a critical role in the concentric organization of OS collicular neurons, whereas the spatial distribution of retinal DS neurons does not account for the organization of DS neurons in the colliculus.

## Results

### Single-bouton imaging reveals the local organization of OS and DS ganglion cell boutons

To investigate the OS and DS tuning of retinal boutons, we measured neuronal calcium responses to a dark bar moving perpendicular to its long edge using two-photon microscopy in awake head-fixed mice (Fig 1C-E). For each responding bouton, an orientation and direction selective index (OSI and DSI) and corresponding p-values (pOSI and pDSI) were calculated. A bouton was deemed orientation and/or direction selective if the relevant selectivity index was greater than 0.1, and corresponding p-value less than 0.01. Eighteen percent of responding boutons were classified as exclusively DS, ten percent were exclusively OS and a small fraction (1.5%) were both (Fig 1G-H), where the orientation selectivity of DS boutons was systematically higher than the direction selectivity of OS boutons (Fig 1I; mean ± SD, 0.17 ± 0.09 and 0.09 ± 0.1, respectively). As a large fraction of boutons were either OS or DS but not both (Fig 1H), OS and DS boutons appear to be largely independent populations, consistent with known response properties of identified ganglion cell populations in the retina ^19,22–24^. Across the depth of the colliculus, we found that OS and DS boutons are distributed differently within the retino-recipient layers. DS boutons were found to dominate the most superficial layers, while OS boutons were more evenly distributed with a peak density at a depth of ∼100 μm (Fig 1K). This is consistent with previous physiological measurements of OS and DS neurons within the colliculus ^10,12,25^, and the anatomical distribution of the axons of genetically identified ganglion cells within the colliculus ^26–29^.We have restricted the rest of the analysis to boutons that were determined to be either OS or DS, both not both (Fig 1H).

**Figure 1:**
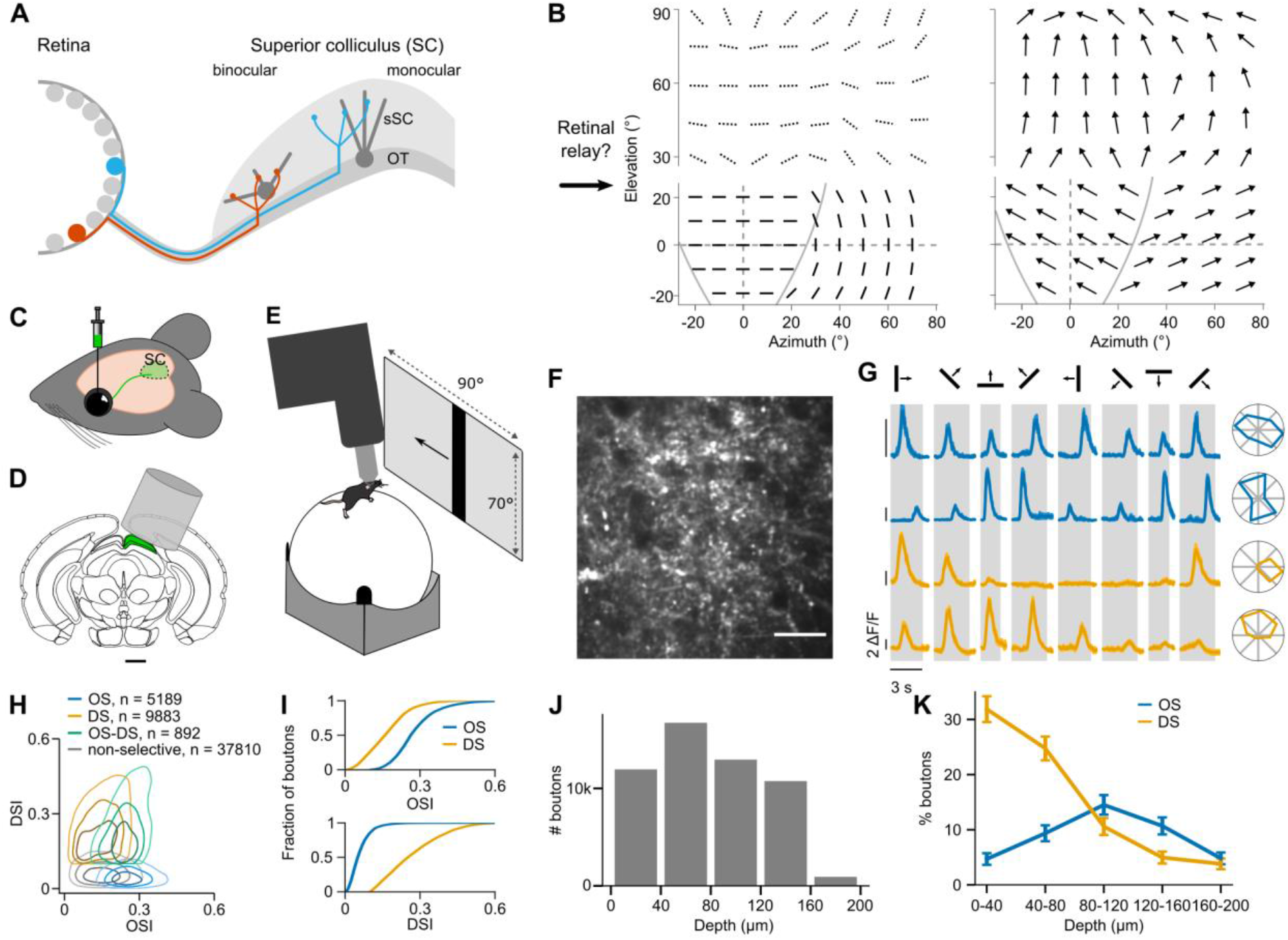
Orientation and direction-selective boutons cluster at different depths of the colliculus. **A**. Schematic of retinotopic innervation of the colliculus by retinal ganglion cells. Dark gray - collicular neurons, red - ganglion cell sampling the binocular zone, blue - ganglion cell sampling the monocular zone. **B**. Visual maps of preferred orientation (left) and direction (right) of neurons in the mouse colliculus. Dashed lines indicate the putative preferred orientation inferred from Li et al., 2020 (*13*). **C**. Intraocular virus injection to label retinal axon boutons. **D**. Implantation of a cranial window on top of the colliculus. **E**. Schematic of the in-vivo recording setup. **F**. Representative field of view of ganglion cell boutons recordings in the colliculus. Scale bar is 20 μm. **G**. Example responses of two OS (blue) and two DS boutons (yellow). **H**. DSI-OSI distributions of OS (blue, OSI > 0.1, pOSI < 0.01, not DS), DS (yellow, DSI > 0.1, pDSI < 0.01, not OS) and OS-DS (green, OSI and DSI > 0.1, pOSI and pDSI < 0.01), and non-selective boutons (gray, not OS nor DS). Contour lines correspond to 25^th^, 50^th^ and 75^th^ percentile. **I**. Cumulative density of OSI values (top) and DSI values (bottom) of OS and DS boutons. **J**. Depth distribution of all recorded responding boutons. **K**. Percentage of OS and DS boutons for each depth bin (mean ± SD). Data from 9 mice, 118 fields of view and 53774 boutons.

To determine whether OS and DS boutons locally cluster based on their orientation and direction preferences in the colliculus, we compared the difference in preferred orientation for pairs of either OS (Fig 2A-D) or DS (Fig 2E-H) boutons. This was done both by looking horizontally (within the same plane) or vertically (within a column). Horizontally, OS boutons showed highly homogeneous preferred orientations within local patches (separation < 90 μm; median [interquartile range], 5 [2-9]°; Fig 2A-C). Across depth, the difference in preferred orientation was also small when depth separation was less than 90 μm (median [interquartile range], 5 [2-10]°, Fig 2D). For more distant pairs, the distribution had two peaks: one close to 0° and the other near 90° (peak 1: median [interquartile range], 8 [4-14]°; peak 2: median [interquartile range], 75 ± [65-83]°, Fig 2D). While in some retinotopic locations, preferred orientations remain homogenous across the sampled depth (Fig 2A), in other locations, preferred orientations rotate by ∼90° between the most superficial and deepest retinorecipient layers (Fig 2B). In contrast, DS boutons within a local patch (∼90 μm) sampled all four cardinal directions (Fig 2E-F) with the distribution of DS pairs often showing two or three peaks at around 0°, 90° and 180° of angular difference (Fig 2E-F). However, boutons located within 20 μm of each other are more likely to prefer the same motion direction (median [interquartile range], 51 [15, 103]°, Fig 2G). Across depth, boutons had the same probability of preferring the same, an opposite or an orthogonal direction (Fig 2H). The data presented above is broadly consistent with the reported local clustering of OS and DS neurons in the colliculus. The similarity is stronger for OS neurons in the colliculus that have been reported to share the same orientation preference within patches ∼100 μm across that span the depth of the vertical thickness of the retinorecipient layers ^10–12^. DS neurons have been reported to form larger clusters of ∼500 μm where neurons prefer the same direction of motion ^11,13^. Within these larger clusters, in particular near the surface of the colliculus, it is common to observe all preferred directions in a local anatomical patch ^11,13,25^. This diversity of preferred directions is common in our recordings of retinal boutons (Fig 2). Together this data highlight the similarity of local clustering of OS collicular neurons and the bouton of OS retinal ganglion cells in the colliculus.

**Figure 2:**
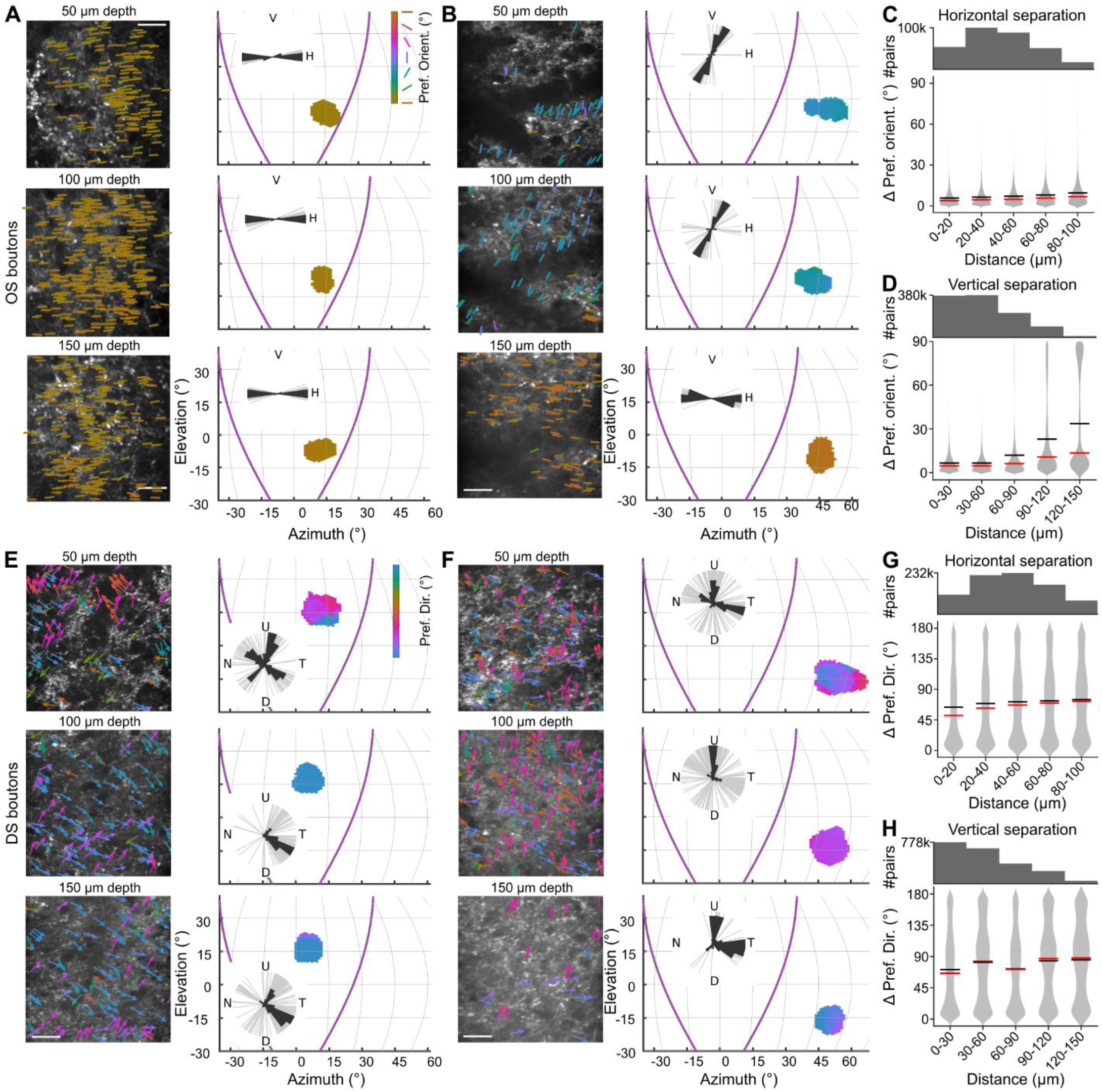
Preferred orientations of OS boutons are locally homogeneous, while DS boutons sample all four cardinal directions. **A-B**. Example columns of OS boutons of the binocular (A) and monocular (B) zone. **Left**. Field of view at 50, 100 and 150 μm depth, perpendicular to the surface of the colliculus. Colored lines indicate preferred orientation of OS boutons. Scale bar is 20 μm. **Right**. Mapping of preferred orientations into visual space. Pixel color corresponds to mean preferred orientation in an area (see colormap). Purple lines separate the binocular and monocular zone. Insets show polar histograms of preferred orientations, individual OS boutons in gray, V-vertical, H-horizontal orientation. **C-D**. Distribution of preferred orientation differences between pairs of OS boutons by horizontal separation within the same imaging window (C) or vertical separation within the same column (D). Red and black horizontal lines indicate median and mean, respectively. **E-F**. Two example columns of DS boutons like (A-B) with preferred directions, U-upward, P-posterior, D-downward, A-anterior. **G-H**. Distribution of preferred direction differences between pairs of DS boutons like (C-D). Data from same animals as Fig 1, C: 95 fields of view, 325253 OS pairs, D: 26 columns, 773648 OS pairs, G: 106 fields of view, 821166 DS pairs, H: 27 columns, 1613830 DS pairs.

### The orientation map of OS retinal boutons match the orientation map of collicular neurons

Given the similarity of the local organization of OS retinal boutons within a local patch, we next assessed whether the global organization of OS retinal boutons matched the concentric organization of OS neurons in the colliculus ^10,11^. To do this, the preferred orientation and receptive field location of each imaging window was mapped into retinotopic space (Fig 3A-B). We found that retinal boutons in the binocular zone prefer horizontal orientations, but a variety of both horizontal and vertical orientations in the monocular zone (Fig 3A and B). Anecdotally, the shifts in orientation preferences across the colliculus of retinal boutons match the concentric organization first reported by Ahmadlou and Heimel ^10^.

**Figure 3:**
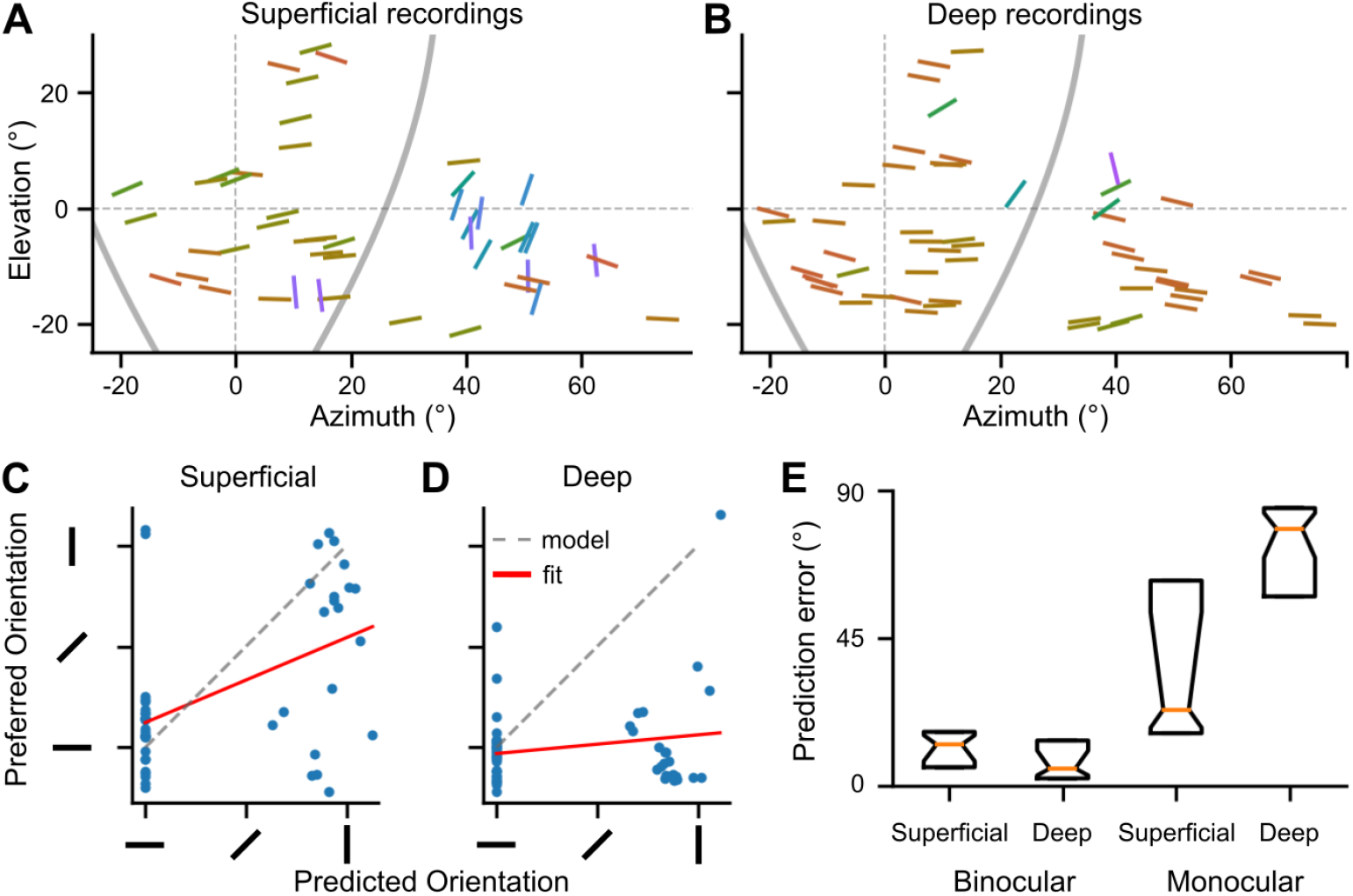
The orientation map of superficial OS boutons aligns with the concentric orientation map of OS SC neurons. **A-B**. Retinotopic maps of mean preferred orientation of each field of view of superficial recordings (A, depth < 60 μm) and deep recordings (B, depth ≥ 60 μm). **C-D**. Preferred orientation of each field of view is compared to its concentric model prediction obtained from receptive field location for superficial (C) and deep recordings (D). Gray dashed line - model, red line - linear fit. **E**. Prediction error between measured preferred orientation and model prediction. Boxplots show median and interquartile range of superficial and deep recordings, divided further into binocular and monocular zones. Notches indicate confidence interval of the median. Data from same animals as Fig 1, 28 upper-binocular, 33 lower-binocular, 20 upper-monocular, 22 lower-monocular fields of view. See also Fig S1

One clear difference between the recordings of OS retinal boutons and OS neurons in the colliculus is how consistent the same angle is encoded across depth. While the same angle is often maintained across the depth of the retinorecipient layers for OS neurons of the colliculus, this is not always the case for the boutons of OS ganglion cells. We observed that boutons at different depths of the colliculus could have different preferred orientations (Fig 2B). To assess the spatial regularity of preferred angle across depth, the colliculus was divided into two regions, the superficial and deep regions, with a border at ∼60 μm. To quantify the similarity of the spatial distribution of preferred orientation between ganglion cell boutons and superior colliculus neurons, we created a prediction model based on previous reports ^10,11^ (Fig 1A and S1).

Specifically, if a recording’s receptive field was located in the monocular zone, the predicted orientation was the concentric angle between the receptive field location and the center point of vision at 0° elevation and azimuth. Whereas, if the recording’s receptive field was in the binocular zone, the predicted orientation was horizontal.

We found that the preferred orientation of boutons in superficial layers matched the predicted preferred orientation of adjacent collicular neurons (fit slope: 0.42, R^2^= 0.23, Fig 3C), those in the deeper layers, below 60 μm, differed from our prediction (fit slope: 0.09, R^2^ = 0.03, Fig 3D). In addition, we computed a prediction error separately for boutons in binocular or monocular zones (Fig 3E, see methods). This confirmed that the orientation map of boutons in the superficial layers fit the collicular orientation map in both the monocular and binocular zones (prediction error in the binocular zone: 13° [6-16]°, in the monocular zone: 23° [16-62]°, median [interquartile range]). Boutons in the deeper layers matched with the modelled collicular orientation map in the binocular zone but not in the monocular zone (prediction error in the binocular zone: 5° [2-14]°, monocular zone: 78° [58-84]°, median [interquartile range]). Together, these observations suggest that the concentric organization of collicular neurons in the monocular zone may be inherited from OS retinal boutons synapsing in the superficial, but not the deep layers of the colliculus.

### There is a general preference for upward and posterior motion across the visual field

The colliculus processes visual motion in a biased manner, where, while more than one preferred direction is often represented at each location, strong preferences exist ^11,25,27,30,31^. In our recording of retinal boutons in the colliculus it was common to find boutons preferring each cardinal direction in all of the ∼100 μm imaging window (Fig 2). However, in most patches there was an overrepresentation of one or two directions, often separated by ∼90 degrees (Fig 2). Therefore, the proportion of DS boutons responding to each cardinal direction was calculated, where each DS bouton was classified into anterior, upward, posterior, or downward groups (Fig 4A, see Methods). We found that upward and posterior preferring DS boutons accounted for ∼80% of all DS boutons recorded in the colliculus (Fig 4A). The strong overrepresentation of upward and posterior motion is similar to the distribution of preferred directions found by Sabbah et al. ^18^ in whole-mount retina recordings, who found ∼72% of ON-OFF DS ganglion cells prefer upward or posterior motion (data replotted in Fig S2B). Analysis by depth showed that DS boutons in the most superficial layers (0 – 60 μm) are more evenly distributed (15% anterior, 30% upwards, 40% posterior and 10% down), while deeper ones (> 150 μm) preferred mainly upward motion (> 60% of boutons) (Fig 4A).

**Figure 4:**
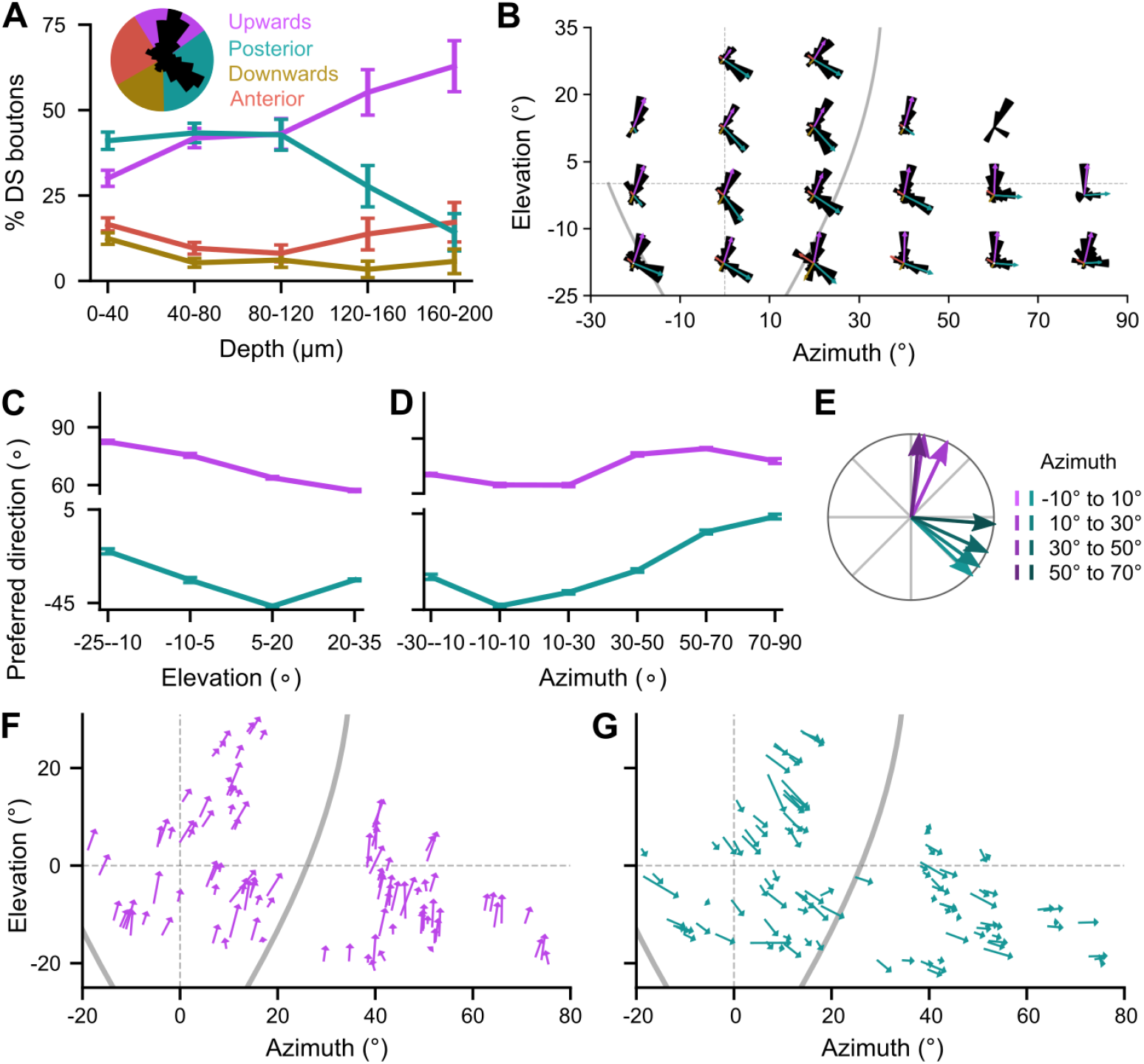
DS boutons show a strong preference for upward and posterior motion aligning to global flow fields. **A**. Depth distribution of DS boutons preferring upward (purple), posterior (teal) and downward (yellow), anterior (red) motion direction (mean ± SD). Inset: Polar distribution of preferred directions of all DS boutons. **B**. Polar distributions of preferred directions at equidistant receptive field locations. Arrows indicate weight and mean preferred direction of each cardinal direction. **C-D**. Mean preferred direction of upward (purple) and posterior motion (teal) preferring boutons by elevation (C) and azimuth (D). **E**. Polar plot indicates the shift of mean directions by azimuth from (D) in polar coordinates. **F-G**. Flow fields of upward (F) and posterior motion preferring boutons (G) determined from mean preferred direction at mean receptive field location of each recording window. Length of arrows indicates percent of anterior and upward motion preferring boutons in each patch. Data from same animals as Fig 1.

To determine if the direction preferences shift across the visual field, each local collicular patch was mapped into retinotopic space. Comparing local preferences across different receptive field locations revealed a dominance for upward and posterior selectivity everywhere in the sampled visual field (Fig 4B), where the mean preferred direction of each cardinal component rotated along both the azimuth and elevation of the receptive field location (Fig 4C-G). Changes between upper and lower visual field were largest for boutons preferring upward motion, shifting from 82° ± 2° to 57° ± 1° (mean ± SD, Fig 4C), whereas changes between the binocular and monocular zones was strongest for boutons preferring posterior motion, shifting from -43° ± 2° to 4° ± 3° (mean ± SD, Fig 4D-E). To highlight the smooth topographic variations of direction preferences across the visual field, we mapped the mean preferred directions of upward and posterior boutons of each recording window. The resulting maps revealed the fine adjustment of upward and posterior preferred directions across the visual field (Fig 4F-G).

These topographic shifts of cardinal preferred directions appear to follow the same organization of DS ganglion cells discovered in the retina ^18^. To compare the map of DS ganglion cell boutons in the colliculus with the map of DS ganglion cells in the retina, we plotted the dataset of Sabbah et al., 2017 ^18^ in the same global visual-field coordinates as the data presented in Fig 4 (Fig S2, see methods). When considering the portion of the visual field sampled by both datasets, we found that the rotation of the mean preferred direction for upward and posterior boutons followed the same trend (Fig S2), indicating that the retinal map of DS ganglion cells appears to be conserved at level of their terminals in the colliculus.

### Ipsilateral retinal inputs lack posterior motion and vertical orientation tuning

Taken together, the above recordings of retinal boutons in the colliculus demonstrate that contralateral retinal inputs match well with the organization of both OS and DS neurons in the lateral visual field, but only to the organization of OS neurons in the binocular visual field (Fig 2-4). As ipsilateral retinal inputs exclusively innervate the binocular zone, we next asked how well ipsilateral OS and DS ganglion cell inputs to the colliculus match their contralateral counterparts. As ipsilateral inputs lie within the deeper portion of the retino-recipient layers of the colliculus ^26^, we recorded the visual response properties of ganglion cells in retinal explants. Ganglion cells were labelled using an AAV expressing GCaMP6s (AAV-hSyn-GCaMP6s) injected into the binocular zone of the colliculus and their visual response properties recorded under a two-photon microscope (Fig 5A-C). Using this technique, we were able to record the visual response properties of ganglion cells projecting either exclusively to the ipsilateral colliculus or a set of neurons that project to binocular zone of the ipsilateral and/or contralateral colliculus (Fig C-D).To identify OS and DS ganglion cells, we stimulated the retinas with drifting gratings moving in one of eight directions inside an aperture centered on the recording window (Fig 5B-C). DS and OS cells were classified based on their selective indexes (see methods). Recording sites were located within the retina’s binocular zone (Fig 5D).

**Figure 5:**
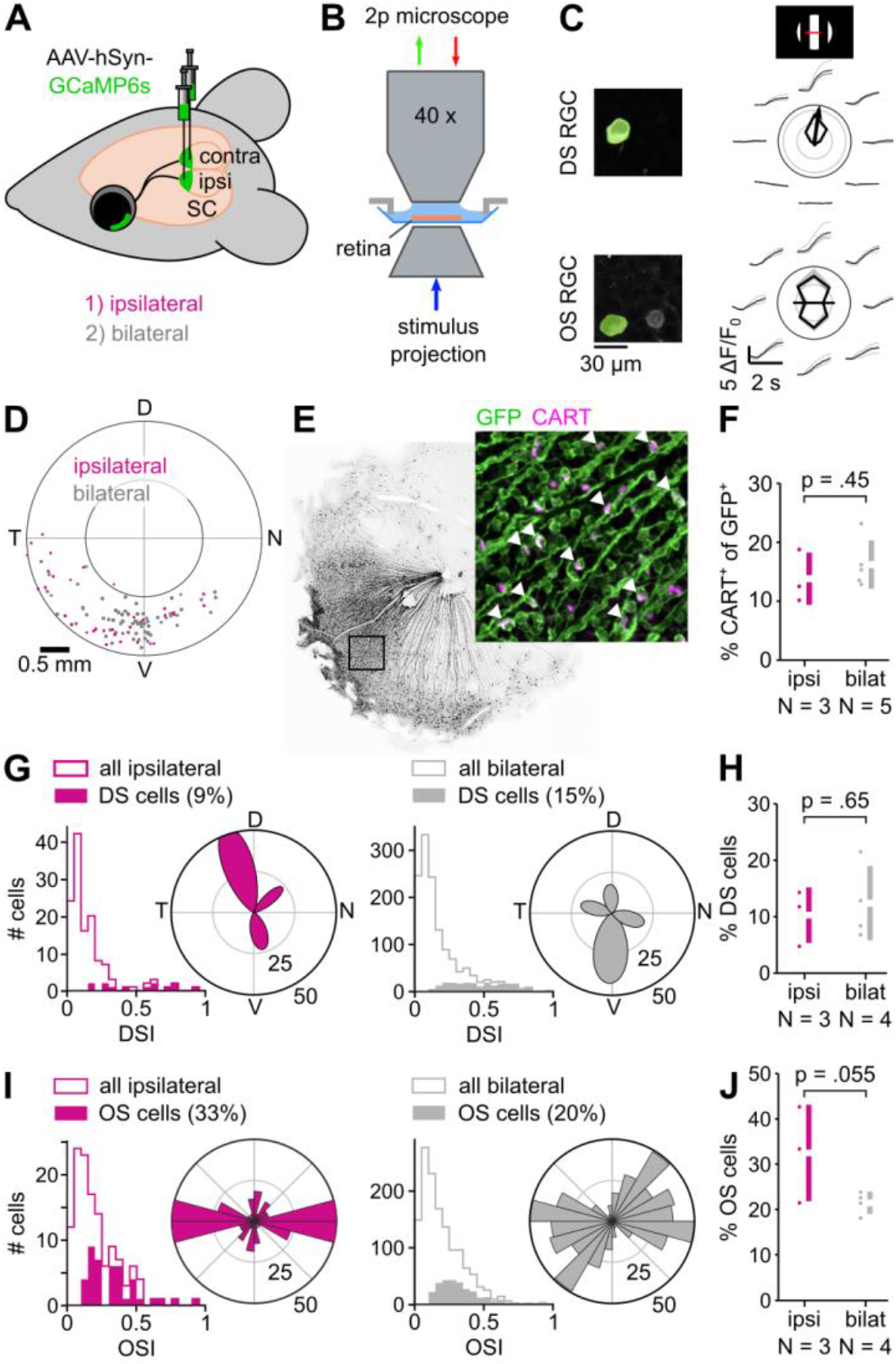
Ipsilateral projections of OS and DS ganglion cells lack posterior motion and vertical orientation tuning, respectively. **A**. Labelling of retinal projections by ipsilateral and bilateral colliculus injections. **B**. Ex-vivo retina calcium imaging and stimulation with drifting gratings. **C**. Example calcium traces of a DS (top) and OS (bottom) ganglion cell. **D**. Recording positions in retinal coordinates. **E**. Retina of bilaterally injected superior colliculus stained for GFP (green) and CART (magenta). Arrows point to double-labelled cells. **F**. Percent of CART+ GFP+ cells from ipsilaterally and bilaterally injected colliculi. Bars: mean ± SD, dots: individual retinas. **G**. Distributions of DSI and preferred directions of DS ganglion cells (DSI > 0.1, pDSI < 0.01) from ipsilateral (left) and bilateral (right) injections. **H**. Percent DS ganglion cells in ipsilateral and bilateral populations. **I-J**. Distributions of OSI and preferred orientations of OS ganglion cells (OSI > 0.1, pOSI < 0.01) (I) as in (G) and percent OS cells (J) as in (H). P-values from permutation t-test of the mean difference. Ipsilateral: N = 3 retinas, 131 ganglion cells; bilateral: N = 4 retinas, 1381 ganglion cells.

First, we consider DS ganglion cells. We estimated the number of DS ganglion cells projecting to the binocular zone by staining each retina after recording for CART, a marker for ON-OFF DS ganglion cells, and GFP (Fig 5E). The percentages of CART positive cells were similar for both exclusively ipsilaterally projecting and mixed ipsi- and contralaterally projecting sets of neurons (ipsilateral: 13.8 ± 3.6 % and bilateral: 16.2 ± 3.7 %, p = 0.45, permutation test, Fig 5F). This proportion was similar to what we found when classifying neurons based on their visual response properties (ipsilateral: 10.3 ± 4.0 % and bilateral: 12.4 ± 5.7 %, p = 0.65, permutation test, Fig 5G-H). To identify a potential bias in the preferred directions of DS inputs, preferred directions were grouped either into three (ipsilateral) or four groups (bilateral, see methods). The number of clusters was decided based on the alignment of the average cluster direction with the four cardinal directions. In the ipsilateral eye, DS cells preferred equally nasal, ventral and dorsal motions in the retina but we did not observe DS cells with a preference for temporal motion (Fig 5G). In contrast, retinas with neurons projecting to both the ipsi- and contralateral colliculus contained DS ganglion cells preferring all four cardinal directions (Fig 5G).

Second, we consider OS ganglion cells. We found that OS inputs from the contralateral eye aligned with the properties of OS neurons in the binocular zone of the colliculus (Fig 3), suggesting that collicular neurons inherit their response preferences from the retina. Here, we examined the OS properties of ipsilaterally projecting ganglion cells (Fig 5). We found a tendency for a higher ratio of OS ganglion cells in ipsilaterally projections compared to mixed set of ipsi- and contralateral projecting ganglion cells (ipsilateral: 32.5 ± 8.7 % and bilateral: 21.4 ± 2.1 %, p = 0.055, permutation test, Fig 5I and J and Fig 1I). Notably, ipsilaterally projecting OS ganglion cells showed strong preference for horizontal orientations (Fig 5I). This aligns well with the orientation preference of contralateral projecting OS retinal boutons and collicular neurons in the binocular zone.

### OS boutons show similar retinotopy as non-selective boutons

The local clustering of OS retinal boutons with the same response properties within the colliculus suggests one of two wiring models. First, ganglion cells send axons to positions in the colliculus that don’t match their corresponding position in the retina, resulting in convergence of similar response properties (Fig 6A). Second, some ganglion cells do not innervate the colliculus, resulting in patches of similar response properties as a consequence of subtraction (Fig 6B). In the case of OS ganglion cells, vertical and horizontal OS ganglion cells appear to uniformly map the retina ^19^. If OS ganglion cells uniformly innervated the colliculus in a precise retinotopic manner ^32,33^, we would expect to find vertical and horizontal preferring OS boutons evenly distributed across the colliculus. Instead, we see local clustering with an absence of vertical OS retinal boutons in the binocular visual field of the colliculus. Therefore, OS ganglion cells might innervate the colliculus in a region-specific fashion. To test if the local clustering of OS boutons by preferred angle of orientation is a consequence of regional convergence or avoidance (Fig 6), the spread in receptive field centers between OS and nonselective boutons was compared (Fig 6C and D). We found that the receptive fields of OS boutons was contained within the distribution of receptive fields of non-selective boutons (Fig 6C). To quantify this trend across all recorded fields of view, we computed the differences between the median and interquartile range (IQR) of RF centers for OS and non-selective boutons (Fig 6D). We found that the difference in median RF centers was close to zero for both azimuth and elevation (azimuth: -0.01 ± [- 0.89, 1.18] °, elevation: -0.31 ± [-1.3, 0.63] °. median ± [IQR]), indicating that OS and non-selective boutons in the same field of view come from ganglion cells at the same retinal location. Furthermore, the spread of non-selective bouton receptive-field centers was greater than that of OS boutons (IQR non-selective – IQR OS: azimuth: 0.89 ± [0.12, 2.1] °, elevation: 1.1 ± [0.14, 2.1] °. median ± [IQR]). Taken together, these results suggest that the patch-like organization of OS ganglion cell boutons in the colliculus, and the absence of vertical orientation is likely the consequence of OS ganglion cells not innervating specific regions of the colliculus.

**Figure 6:**
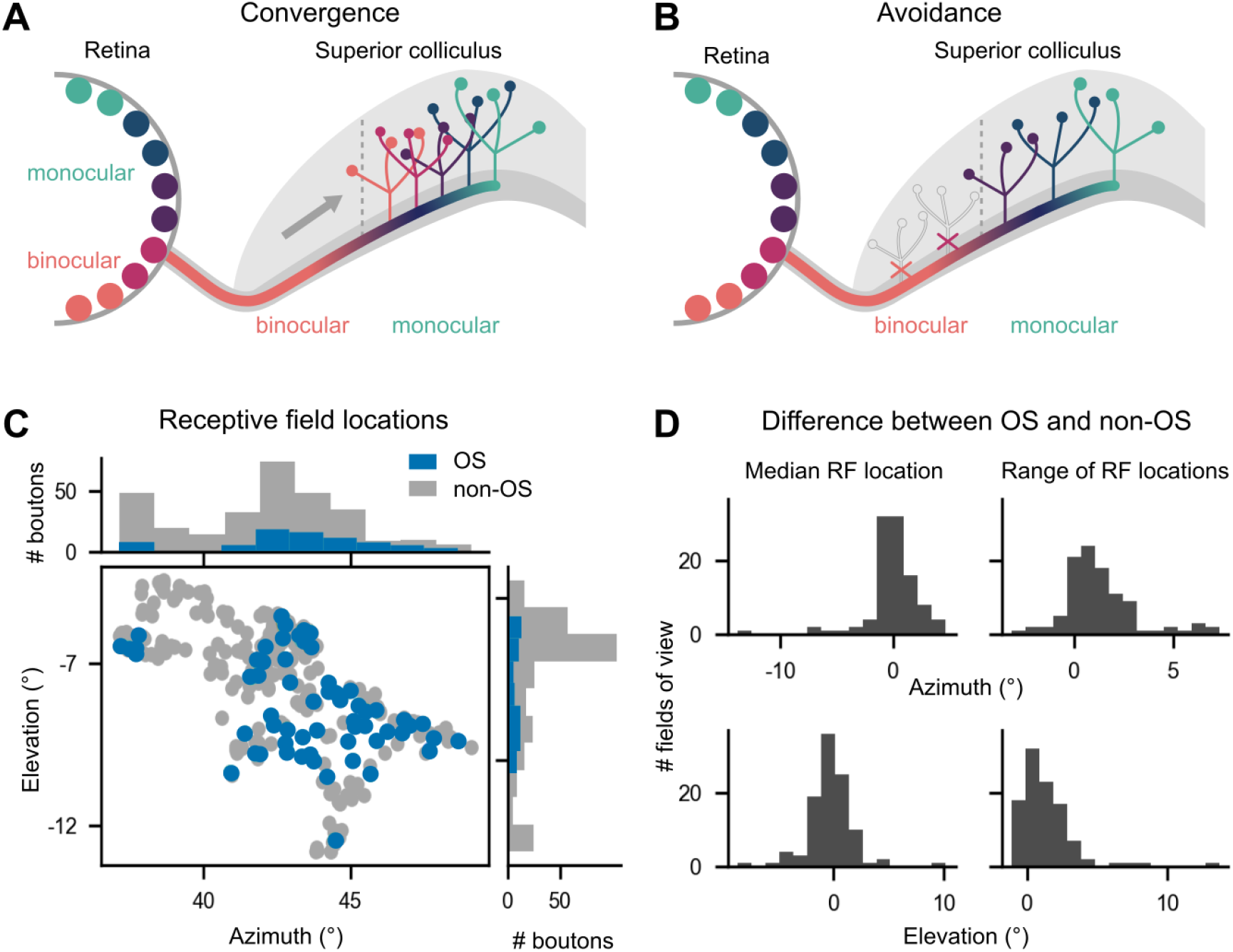
OS boutons show a similar retinotopic mapping as non-OS boutons. **A-B**. Region-specific innervation of retinal boutons by convergence (A) or avoidance (B). **C**. Distribution of receptive field (RF) locations of OS boutons (blue) and non-OS boutons (gray) in a single field of view (Fig. 2B, middle). **D**. Comparison of median (left) and interquartile range (right) of RF locations between OS and non-OS ganglion cell boutons for each field of view. Distributions of differences for azimuth (top) and elevation (bottom). A positive difference indicates that the median or range of non-OS boutons is bigger than the one of OS boutons. Data from same animals as Fig 1.

## Discussion

Here, we investigated whether the retinotopic maps of orientation and direction observed in the superior colliculus are directly inherited from the retina. We made three key observations. First, both OS and DS ganglion cell boutons tend to locally cluster by preferred angle within the colliculus (Fig 2). Second, while the orientation map of retinal boutons was consistent with the orientation map of collicular neurons (Fig 3), direction maps of DS boutons did not display regional biases (Fig 4). Instead, DS boutons showed a global bias for upward and posterior motion preference, reproducing the spatial organization found in the retina. Third, we found that orientation and direction selective boutons formed two distinct groups, where the distribution patterns differed near the surface compared to boutons lying deeper in retino-recipient layers of the colliculus (Fig 1-2). Together this work suggests that while the global pattern of orientation found in the colliculus is consistent with the organization of retinal inputs, the organization of direction is not (Fig 7).

**Figure 7:**
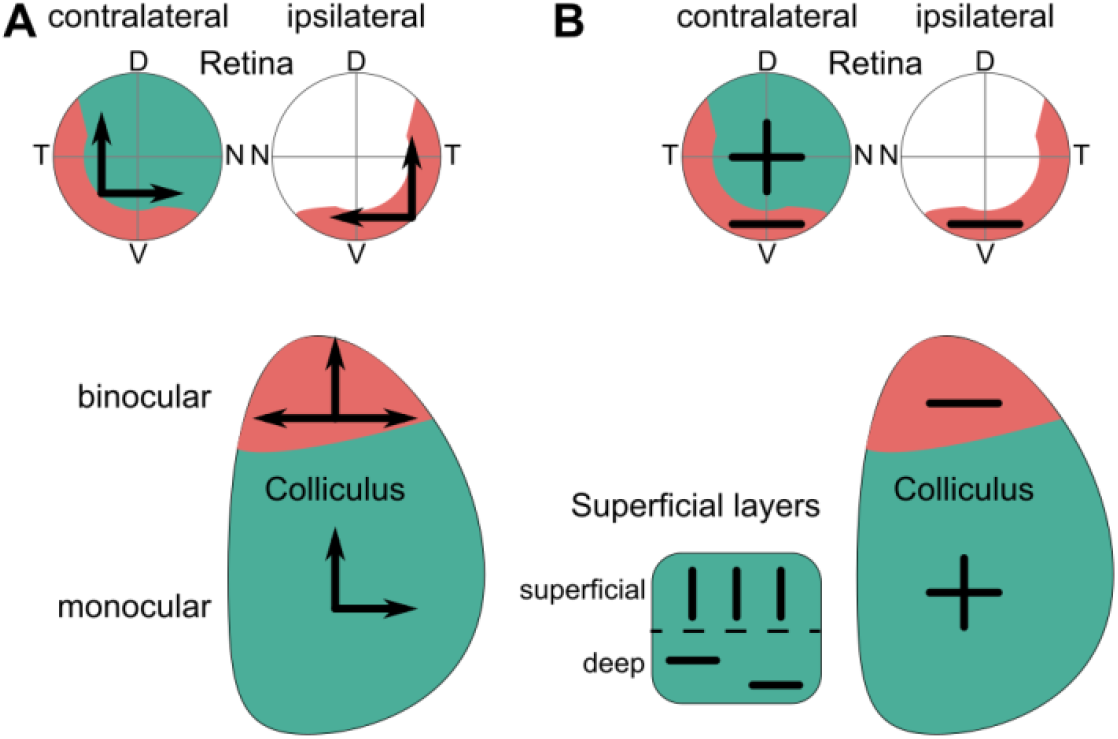
Summary of retinal inheritance of orientation and direction maps. **A**. DS inputs from the contralateral eye show global bias for upward and posterior motion, resulting in a bias for upward and posterior motion in collicular neurons of the monocular zone. DS inputs from the ipsilateral eye lack posterior motion, potentially contributing to a different bias in collicular neurons of the binocular zone. **B**. OS inputs from the binocular zone of both, the ipsi- and contralateral eye, are tuned to horizontal orientation. In the monocular zone, boutons preferring vertical orientation can be found in the more superficial layers and boutons preferring horizontal orientation in the deeper layers.

The local clustering of neurons in the colliculus with similar orientation and direction preferences has been an interesting recent observation ^10–13^. Recordings from retinal boutons in the colliculus have not, to date, identified regional clustering of cells by response preference ^32–34^. Here we show that both OS and DS retinal inputs show evidence for clustering based on their preferred orientation or direction (Fig 2). The pattern of local clustering of OS retinal boutons (Fig 2-3) matched well with the concentric arrangement of OS neurons around the center of vision observed in the colliculus ^10^. However, two global patterns of DS neurons in the colliculus ^11,13^ were not matched by the spatial distribution of DS retinal boutons (Fig 2 and 4). Instead, particularly in the most superficial layers of the colliculus it was common to observe boutons of each of the 4 preferred directions. This more closely matches the observations of Inayat et al., 2015 ^25^, who reported no local clustering of DS neurons at the surface of the colliculus.

### Layer-specific organization of OS and DS inputs

Our recordings of OS and DS retinal boutons revealed differences between the groups of boutons in the most superficial layers and those that lay deeper in the colliculus (Fig 2 and 3). These differences were more pronounced for DS retinal boutons, where it was common to find a decrease in the number of preferred directions at deeper depths, but representation of all 4 cardinal directions near the surface (Fig 2). Consistent with our recording, three of the ON-OFF DS ganglion cell subtypes are known to stratify in the most superficial layers of the colliculus ^24,27,29^. In addition, the upward motion preferring ON-OFF DS ganglion cells show the broadest and deepest innervations patterns of the genetically identified ON-OFF DS ganglion cells, stratifying at depths down to 150 μm in accordance with both our observations of more upward motion preferring boutons in the deeper layers (Fig 5), and previous recordings using electrophysiology that have difficultly recording neurons at the surface ^30,31^.

Unlike DS retinal boutons, the preferred orientation of OS retinal boutons was found to be relatively consistent across depth, with a few observations of differences at depths of more than 90 μm (Fig 2). Critically, while in the superficial layers (top 90 μm) we recorded OS boutons with a variety of preferred orientations, deeper layers were dominated by OS boutons that preferred horizontal orientations (Fig 2). This is consistent with our observation that OS ipsilaterally projecting ganglion cells, known to stratify in the deep layers, are selective for horizontal orientations and in the binocular zone it has been observed that there is a clear contribution to horizontal preference from both the ipsi- and contralateral eye ^35^. These observations are consistent with the depth profiles of recorded OS neuron in the colliculus and the stratification patterns of vertically tuned OS neurons (JAMB^+^) that stratify in the upper layer of the superficial superior colliculus ^22,27^. Due to the absence of a mouse line that labels horizontal orientation preferring ganglion cells, we can only speculate about their stratification patterns.

### Evidence of region-specific inputs of OS ganglion cells to the superior colliculus

Previous studies have observed topographic variations in the strength of a specific retinal cell-type’s to the colliculus of cat’s ^36,37^. Here we observed that OS ganglion cells that preferred vertical orientations were absent in the binocular zone but are present in a sub-region of the sampled monocular zone. This is particularly intriguing given that OS ganglion cells preferring vertical orientations have been recorded across all retinotopic locations in the retina ^19,22^. However, our results agree with anatomical evidence of region-specific wiring of genetically identified ganglion cell types into the superior colliculus ^23,26,27^. For example, despite a uniform coverage of the retina the axonal arborizations of JAMB^+^ ganglion cells in the colliculus are patchy, with holes of up to 500 μm ^27^. These holes may account for the missing vertical orientation in the binocular zone and could explain the mismatch between the coverage of the retina by OS ganglion cells tuned to vertical orientation and the coverage of the colliculus by their boutons. The proposed avoidance model (Fig 6) could be implemented either by axons never innervating specific patches of the colliculus, or by pruning of the axon terminals during development, perhaps guided by visual experience.

### Collicular direction maps are only partially formed by input from DS ganglion cells

Direction selectivity in collicular neurons requires input from DS ganglion cells ^20^. While, each DS ganglion cell subtype covers the retina homogeneously, a global bias for posterior and upward motion does exist ^18^. This global bias resembles the retinotopic direction map of DS boutons in the colliculus observed by us and others ^34^. Together, this suggests a one-to-one feedforward transmission of the retinal DS inputs without region-specific wiring. However, these feedforward maps cannot account for the regional specialization observed in collicular neurons ^11,13^. The creation of these regional specializations likely requires another set of inputs, which could include external inputs from the visual thalamus and cortex ^38,39^, or local circuit computations mediated by collicular interneurons ^40–43^. For example, the colliculus receives inhibitory input from the vLGN which in turn receives input from TRHR^+^ posterior motion preferring DS ganglion cells ^24,44,45^. The posterior motion preferring cells of the vLGN could then inhibit untuned cells in the colliculus to become anterior motion preferring.

### Analogous models of colliculus orientation and direction

Here, we provide a template of how orientation and direction maps of colliculus neurons could be formed based on previous reports ^10–13^. First, direction maps have been reported to have a bias for upward and anterior motion preference in the binocular zone and for upward and posterior motion in the monocular zone. The observed bias for upward motion could be a simple consequence of the strong bias of DS retinal inputs for upward motion (Fig 2). Second, it has been reported that in the anterior visual field, orientation maps follow concentric circles ^10^. The spatial distribution of OS retinal boutons, observed here, is consistent with the concentric maps reported by Ahmadlou and Heimel (2015) ^10^. In addition, our data demonstrate an almost exclusive representation of horizontal orientation in the binocular zone as also observed by Russell et al., (2022) ^35^. Hence, our proposed scheme seems a valid approximation of the orientation map in the anterior colliculus.

### Feedforward transfer of regional orientation preference in circuits driving innate behavior

We have shown that the map of preferred orientation of OS ganglion cell boutons coincides with the same map previously reported for superior colliculus’ neurons. In addition, we found that the orientation tuned input coming from both eyes in the binocular zone aligns with the horizontal orientation observed in the rostral colliculus. It is, thus, tempting to explain the organization of the collicular map with a simple feedforward model of orientation responses from the retina to the colliculus. This is a clear demonstration of how region-specific wiring of one visual area to the next can contribute to the creation of visual feature maps, independent of the topographic map in the upstream area. The downstream continuation of these topographic maps is suggested by the distinct wiring of the binocular and monocular zone of the colliculus to the parabigeminal nucleus and lateral posterior nucleus of the thalamus ^46,47^. This might aid innate orienting behaviors similar to toads ^48^ where horizontal orientation triggers approach and vertical orientation avoidance. As mice keep their prey within the binocular zone during hunting ^49^, the preference for horizontal orientation in the binocular zone might aid hunting by detecting horizontally oriented prey.

## Acknowledgments

This work was supported by the FWO (G094616N to KF, G091719N to KF, 1205421N to NKK) and the European Union’s Horizon 2020 research and innovation program under the Marie Sklodowska-Curie grant agreement no. 796102 and 894697 to NKK and DdM, respectively. Thanks to Adrien Philippon for developing the air-cushioned Styrofoam ball setup.

## Author contributions

Conceptualization, DdM, NKK, KF; Experimental setup and development, DdM, NKK, CL; experiments, in-vivo imaging, DdM, NKK, retina imaging CL; analysis, DdM, NKK; software, DdM, NKK; writing & editing, DdM, NKK, KF.

## Materials and Methods

### Resources Table

| Reagent / Resource | Designation | Source or reference | Identifiers |
| --- | --- | --- | --- |
| Genetic reagent (Mus musculus) | C57BL/6J | The Jackson laboratory | JAX: #000664 |
| Antibody | anti-GFP (chicken, polyclonal, 1:500) | Thermo Fisher Scientific | Cat# A-10262;<br>RRID: AB_2534023 |
| Antibody | Alexa 488 donkey anti-chicken 1:500 | Immuno-Jackson | Cat# 703-545-155<br>RRID: AB_2340375 |
| Antibody | anti-CART (rabbit, polyclonal, 1:500) | Phoenix Pharmaceuticals | H-003-62<br>RRID: AB_2313614 |
| Antibody | DyLight 405 donkey anti-rabbit 1:400 | Immuno-Jackson | Cat# 715-475-150<br>RRID: AB_2340839 |
| Antibody | DAPI (1:2000) | Roche | Cat# 10276236001 |
| Tracer | Lipophilic tracers DiD | Thermo Fisher Scientific | Cat# D7776, D7757, D275 |
| Virus | AAV1-hSyn-Flex-GCaMP6s | addgene | #105714 |
| Chemicals, peptides, and recombinant proteins |  |  |  |
| Reagent | 10x PBS | VWR | Cat# 437117K |
| Reagent | 1x PBS | VWR | Cat# 444057Y |
| Reagent | Histofix 4% | Roche | Cat# P087.5 |
| Reagent | Normal Donkey Serum | Millipore | Cat# 30-100ML |
| Reagent | 10% Bovine Albumin | Sigma | Cat# SRE0036-250ML |
| Reagent | DMEM, high glucose | Thermo Fisher Scientific | Cat# 41965062 |
| Reagent | Trypsin 0.05% | Thermo Fisher Scientific | Cat# 25300054 |
| Reagent | Fetal Bovine Serum (FBS) | Thermo Fisher Scientific | Cat# 10270106 |
| Reagent | 2,2'-thiodiethanol (TDE) | Sigma | Cat# 166782-500G |
| Reagent | ProLong® Gold Antifade Mounting Medium | Thermo Fisher Scientific | Cat# P36934 |
| Chemical compound, drug | Sodium Azide (NaN<sub>3</sub>) | Sigma | Cat# S2002-100G |
| Chemical compound, drug | Triton X-100 | Sigma | Cat# S8875 |
| Chemical compound, drug | Clozapine N-oxide (CNO) | Sigma | C0832 |
| Chemical compound, drug | Sodium Chloride (NaCl) | Sigma | Cat# S7653-250G |
| Chemical compound, drug | Potassium Chloride (KCl) | Sigma | Cat# P5405-25G |
| Chemical compound, drug | Calcium Chloride (CaCl<sub>2</sub>) | Sigma | Cat# C5670-100G |
| Chemical compound, drug | Magnesium Chloride (MgCl<sub>2</sub>) | Sigma | Cat# 4880 |
| Chemical compound, drug | D-glucose (Dextrose) | Sigma | Cat# D9434-250G |
| Chemical compound, drug | Sodium phosphate monobasic (NaH<sub>2</sub>PO<sub>4</sub>) | Sigma | Cat# S5011 |
| Chemical compound, drug | Sodium Hydroxide (NaOH) | Sigma | Cat# 655104-500G |
| Chemical compound, drug | Sodium bicarbonate (NaHCO<sub>3</sub>) | Sigma | Cat# S8875-1KG |
| Software and algorithms |  |  |  |
| Software | Fiji | Schindelin et al. 2012 | RRID: SCR_002285 |
| Software | MATLAB | Mathworks | RRID: SCR_001622 |
| Software | Zen lite | Zeiss |  |
| Software | PYTHON 3.4 | Python Software Foundation | www.python.org |
| Software | CaImAn 1.7.1 | Flatiron Institute | https://github.com/flatironinstitute/CaImAn |
| Software | GNU Octave | Free Software Foundation | www.gnu.org/software/octave |
| Software | Psychophysics Toolbox | Psychtoolbox | http://psychtoolbox.org |
| Software | WaveSurfer (version: 0.918) | Janelia Research Campus | http://wavesurfer.janelia.org/ |
| Software | ScanImage | Vidrio Technologies | http://scanimage.vidriotechnologies.com |
| Other |  |  |  |
| Other | Rapid Flow Filters 0.2 µm pore size | VWR | Cat# 514-0027 |
| Other | Wiretrol® II capillary micropipettes | Drumond Scientific | Cat# 5-000-2005 |
| Other | Borosilicate glass | Sutter Instrument | Cat# BF100-20-10 |
| Other | Glass cylinders, Ø 3 mm |  |  |
| Other | Round borosilicate cover glasses, Ø 3 mm, 0.1 mm thickness |  |  |
| Other | UV-curing optical adhesive | Norland | 61 |
| Other | Metabond dental cement | Crown & Bridge |  |
| Other | Super-Bond C&B kit for self-curing adhesive resin cement | Sun Medical |  |
| Other | Laser-Based Micropipette Puller | Sutter Instrument | Cat# P-2000 |
| Other | Small Animal Stereotaxic Workstation | Narishige | Cat# SR-5N |
| Other | Stereotaxic Micromanipulator | Narishige | Cat# SM-15R |
| Other | Hydraulic Oil Micromanipulator | Narishige | Cat# MO-10 |
| Other | Oil Microinjector | Narishige | Cat# IM-9B |
| Other | Two-photon microscope | Scientifica | Serial# 14200 |
| Other | 780 nm LED light source | Thorlabs | Cat# M780L3 |
| Other | Zeiss LSM 710 confocal microscope | Zeiss | Cat# LSM710 |
| Other | Mai Tai HP Ti:Sapphire laser | Spectra-Physics |  |
| Other | B-Scope 2p microscope | Thorlabs |  |

### Mice

All experimental procedures were approved by the Ethical Committee for Animal Experimentation (ECD) of the KU Leuven and followed the European Communities Guidelines on the Care and Use of Laboratory Animals (014-2018/EEC, 166-2018/EEC). Both male and female adult (1-3 months old) C57Bl6J mice were used in our experiments. Mice were kept on a 12-h light-dark cycle (lights on at 7:00), and sterilized food pellets and water were provided *ad libitum*.

### Virus strategy

#### Labelling of ganglion cell boutons

2 ul of AAV-hSyn-GCaMP6s coding for the fluorescent calcium indicator GCaMP6s were intraocularly injected to the right eye of C57Bl6J mice of 6 weeks age.

#### Labelling of ganglion cells projecting to colliculus

100 nl of AAV-hSyn-GCaMP6s were injected into the binocular zone of the left colliculus.

### Stereotaxic surgery

#### Eye and brain injections

Animals were quickly anesthetized with Isoflurane (Iso-vet 1000mg/ml) and a mixture of Ketamine and Medetomidine (0.75 mL Ketamine (100 mg/mL) + 1 mL Medetomidine (1 mg/mL) + 8.2 mL Saline). Mice were placed in a stereotaxic workstation (Narishige, SR-5N). We used micropipettes (Wiretrol® II capillary micropipettes, Drumond Scientific, 5-000-2005) with an open tip of around 30 μm and an oil-based hydraulic micromanipulator MO-10 (Narishige) for viurs injections. Dura tear (NOVARTIS, 288/28062-7) was applied to protect the eyes. To label the injection sites, DiD (Thermo, D7757) was used to coat the pipette tip. The injection coordinates for the binocular zone of the colliculus in an 8-week-old mouse with a bregma-lambda distance of 4.2 mm were AP: −3.2; ML: 0.5; DV: 1.3 mm. Following injection, the wound was closed using Vetbond tissue adhesive (3M,1469). After surgery, mice were allowed to recover on top of a heating pad and were provided with soft food and water containing antibiotics (emdotrim, ecuphar, BE-V235523).

#### Cranial window implantation

A cranial window was implanted above the superior colliculus following the eye injection during the same procedure. A metal head post was fixated on the skull, and a circular 4 mm craniotomy was performed above the frontal two-thirds of the left superior colliculus (ML: 0.0 to 4.0 mm, AP: -0.7 to -4.7 mm). The brain above the colliculus was removed and a cannular window of 3 mm diameter and 2 mm height was implanted and fixated with dental cement (de Malmazet, Kühn, and Farrow 2018).

### Immunohistochemistry

#### Retina immunohistochemistry

Dissected retinas were fixed in 4% paraformaldehyde (PFA, Histofix, ROTH, P087.5mm) with 100 mM sucrose for 30 min at 4°C, and washed overnight or longer at 4°C. After washing, retinas were transferred to wells containing 10%, 20%, and 30% sucrose solution and allowed to sink for a minimum of 30 min, 1hr at room temperature and overnight at 4°C, respectively. The next day, freeze-cracking was performed: retinas were frozen on a slide fully covered with 30% sucrose for 5 min on dry ice. The slides were then thawed at room temperature. The freeze-thaw cycle was repeated three times. Retinas were washed 3 times for 10 min each in 1x PBS, followed by incubation with blocking buffer (10% NDS, 1% BSA, 0.5% TritonX-100, 0.02% NaN_3_ in 1x PBS) for at least 1hr at room temperature or overnight. Both primary antibodies and secondary antibodies were diluted with 3% NDS, 1% BSA, 0.5% TritonX-100, 0.02% NaN_3_ in 1x PBS. Retinas were first incubated with primary antibody solution for 5-7 days under constant gentle shaking at room temperature. Then, retinas were washed three times, 15 min each, in 1x PBS with 0.5% TritonX-100 before being transferred into the secondary antibody solution. Primary antibodies used were chicken anti-GFP (Invitrogen, A-10262, 1:500) and rabbit anti-CART (Phoenix, H-003-62,1:500). Secondary antibodies were Alexa488 donkey anti-chicken (ImmunoJackson, 703-545-155, 1:500) and DyLight 405 donkey anti-rabbit (ImmunoJackson, 715-475-150, 1:200). Retinas were covered with mounting medium and a glass coverslip.

#### Brain immunohistochemistry

Extracted brains were post-fixed in 4% PFA overnight at 4°C. Vibratome coronal sections (100 μm) were collected in 1x PBS and were incubated in blocking buffer (1x PBS, 0.3% Triton X-100, 10% Donkey serum) at room temperature for at least 1 hour or overnight. Then brain slices were incubated with primary antibodies solution for 2-3 days at 4°C with shaking. Slices were later washed 3 times for 10 min each in 1x PBS with 0.3% TritonX-100 and incubated in secondary antibody solution for 2-3 days at 4°C. Primary and secondary antibody were chicken anti-GFP (1:1000) and Alexa488 donkey anti-chicken (1:800), respectively. Nuclei were stained with DAPI (1:1000) together with the secondary antibody solution. Sections were then again washed 3 times for 10 min in 1x PBS with 0.3% TritonX-100 and 1 time in 1x PBS, covered with mounting medium and a glass coverslip. PBS were prepared with 0.02% NaN3.

### Confocal microscopy

Confocal microscopy was performed on a Zeiss LSM 710 microscope. Overview images of the retina and brain were obtained with a 10x (plan-APOCHROMAT 0.45 NA, Zeiss) objective. The following settings were used for imaging the whole-mount retina: zoom 0.7, 4 × 4 tiles with 0% to 15% overlap, 2.37 μm/pixel resolution. For single ganglion cell scanning, we used a 63x (plan-APOCHROMAT 1.4 NA, Zeiss) objective. The following settings were used: zoom 0.7, 2 × 2 tiles or more (depending on size and number of cells) with 0% to 15% overlap. This resulted in an XY-resolution of 0.38 μm/pixel and a Z-resolution between 0.25 and 0.35 μm/pixel. The Z-stacks covered approximately 50 μm in depth. The whole-brain images were acquired with the 10x objective with zoom 0.7, multiple tiles with 5% to 15% overlap and a z-stack of 20 um.

### Ex-vivo retina recordings

#### Tissue preparation

Mice were dark-adapted for a minimum of 30 minutes. After cervical dislocation, eyes were gently touched with a soldering iron (Weller, BP650) to label the nasal part of the cornea and then enucleated. Retina isolation was done under deep red illumination in Ringer’s medium (110 mM NaCl, 2.5 mM KCl, 1 mM CaCl2, 1.6 mM MgCl2, 10 mM D-glucose, 22 mM NaHCO3, bubbled with 5% CO2/95% O2, pH 7.4). The retinas were then mounted ganglion cell-side up on filter paper (Millipore, HAWP01300) that had a 3.5 mm wide rectangular aperture in the center, and superfused with Ringer’s medium at 32–36°C in the microscope chamber for the duration of the experiment.

#### Two-photon recording

Fluorescent cells expressing GCaMP6s were targeted for recording using a two-photon microscope (Scientifica) equipped with a Mai Tai HP two-photon laser (Spectra Physics) at 920 nm wavelength. To facilitate targeting, two-photon fluorescent images were overlaid with the IR image acquired through a CCD camera. Infrared light was produced using the light from an LED. For some cells, z-stacks were acquired using ScanImage (Vidrio Technologies). Imaging frames were of 230 x 57 μm^2^, sampled at 6.8 Hz (bidirectional scanning) and 512 x 128 pixels resolution.

#### Visual stimulation

Stimuli were generated with an LCD projector (Samsung, SP F10M) at a refresh rate of 60 Hz, controlled with custom software written in Octave based on Psychtoolbox (Kleiner et al. 2007). The projector produced a light spectrum that ranged from ∼430 nm to ∼670 nm. The power produced by the projector was 300 lux (at full brightness blue screen and with short pass filter) at the retina. A combination of a neutral density filter and a short pass filter (cutoff wavelength = 475 nm) were used to control the stimulus intensity in logarithmic steps. Recordings were performed with filters decreasing the stimulus intensity by 1-2 log units.

Drifting square-wave gratings of 200 μm periodicity and 2 Hz frequency were shown in eight equidistant directions of random order in an aperture of 300 μm diameter on a dark background.

### In-vivo recordings of retinal boutons in the colliculus

Retinal boutons were labelled with a fluorescent calcium indicator by virus injections into the contralateral eye (see Viral Injections). Visual responses were recorded by two-photon calcium imaging through a cranial window in head-fixed, awake mice.

#### Two-photon calcium imaging setup

A commercial two-photon galvo-resonance scanning microscope with a rotating arm (Thorlabs Multiphoton Microscope, B-Scope) was used to image the calcium signals of retinal boutons in the superficial layers of the superior colliculus. Boutons were imaged at different depth, perpendicular to the surface of the colliculus, using a piezo-electric linear actuator (P-665, Physik Instrumente) to move the objective (Nikon 16x, 0.8 NA). Imaging frames were of 100 x 100 μm^2^, sampled at 60 Hz (bidirectional scanning) and 256x256 pixels resolution. GCaMP6s was excited with a laser at 920 nm wavelength (Mai Tai DeepSee, Spectra Physics). A Pockel’s cell was used to shut down the laser during flyback and to adjust its power leading to a nominal laser power of 30-90 mW at the end of the objective. The head-fixed animals were free to run on a custom 3D-printed air-cushioned spherical treadmill (polystyrene white ball Ø 20cm). We recorded the movement of the ball using two motion sensors (Tindie, PMW3360) with an analog readout of the pitch of the ball in direction of the animal’s head. Eye movements of the mice were estimated by recording the left eye (ipsilateral to colliculus) with a lateral camera (Allied vision, mako G-030B + Lens: NAVITAR, HR F1.4/25MM) positioned at 25 cm distance. The microscope was controlled, and the imaging frames acquired using ScanImage 5.4. Running speed and trigger signals of stimulus beginning and end, two-photon imaging frames and camera frames were recorded in WaveSurfer 0.97, yoked to ScanImage.

#### Mouse training

Prior to the first recording session, mice were habituated to human handling for three days. Then they were head-fixed on the air-cushioned spherical treadmill for 5 minutes on the first day, with increasing durations of 10 min, 30 min and 1 h in the consecutive days. Animals usually adapted easily to running on the treadmill.

#### Presentation of Visual Stimuli

Stimuli were presented in grayscale on a curved LCD monitor (Samsung S32E590C, 729 x 425 mm^2^, 2.9 cd/cm^2^ mean luminance, 60 Hz refresh rate), controlled by custom software written in Octave using the Psychtoolbox. The monitor was at 30 cm distance and covered a visual field of 100° azimuth and 70° elevation on the right side of the mouse (contralateral to colliculus). The irradiance at the position of the mouse ranged from 0.15 to 30 lux (black to white screen). Gray values were gamma corrected to yield a linearly perceived change in brightness. The following visual stimulus were shown to the animal (white, grey and black correspond to 100%, 0% and -100% Weber contrast). A TTL pulse was sent at the beginning and the end of each repeat for later alignment.

#### Moving bar

A black bar of 8° width and extended till the edge of the screen moved perpendicular to its orientation in 8 directions (0°, 45°, 90°, 135°, 180°, 225°, 270°, 315°) with a speed of 40°/s across the screen. Each direction was repeated 4 times. The directions were randomized and interleaved by 3 s of grey screen.

### Analysis

All data analysis was performed in either Matlab or Python.

#### Two-photon image registration and ROI extraction

Two-photon imaging frames were registered, and ROIs determined using NNMF with CamAn 1.7.1 (Flatiron Institute) in Python 3.4. The signal decay time for the model was adjusted to GCaMP6s (1.4 s). ΔF/F values of these selected ROIs were used for further analysis. *Bouton and cell selection*. The repeats of visual stimuli and neural responses were aligned by using the simultaneously recorded stimulus start and imaging frame triggers. For each presented stimulus, a quality index of each ROI was calculated. The quality index was calculated from the variance of ΔF/F across time, var(·)_t_, and the mean across stimulus repeats, ⟨·⟩_r_, as follows:

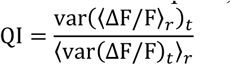

Retinal boutons and cells were included into the analysis if they were visually responsive with a quality index above 0.6.

#### Orientation and direction selectivity index

The orientation and direction selectivity of retinal boutons and ganglion cells was analyzed by calculating the direction selectivity index (DSI) and orientation selectivity index (OSI), preferred directions and preferred orientations from mean peak responses *r*_*θ*_ to each direction *θ* of the moving bar stimulus:

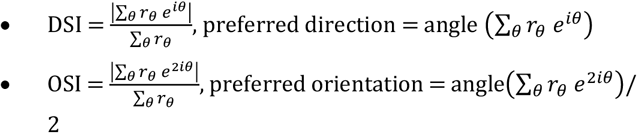

#### Classification of OS and DS ganglion cells and boutons

Ganglion cells and their boutons were considered direction or orientation selective when their DSI or OSI was above 0.1 and the corresponding p-value below 0.01 (99% confidence interval), respectively. The few percent of boutons that were both OS and DS (<3%), were excluded from the analysis of the orientation and direction maps.

#### Receptive field location

Receptive field locations of individual boutons were inferred based on the response delay to the moving bar stimulus, as previously detailed ^11^.

#### DSI and OSI distributions

Contour plots and cumulative density distributions of DSI and OSI values were plotted using the kernel density estimate (KDE) from python seaborn, normalized by cell type (OS, DS, OS-DS, non-selective). The smoothing bandwidth was determined by Scott’s method. Contours correspond to the 25^th^, 50^th^, and 75^th^ percentile.

#### Depth profiles of retinal boutons in the colliculus

To create the depth profile of percentage of OS and DS boutons and of DS preferring each cardinal direction, we binned boutons by depth within 40 μm from 0 to 200 μm depth. To estimate the error in the percentages, we performed the following bootstrapping analysis. For every repeat, 300 boutons from each depth bin were randomly selected with replacement and the percentages previously described were computed. This process was repeated 10000 times, and finally the standard deviations of each percentage across all repeats were computed.

#### Orientation map model comparison

The preferred orientation of OS boutons was compared to a retinotopic model of preferred orientation of colliculus neurons, adapted from Ahmadlou and Heimel 2015. This model derives the preferred orientation of a neuron or bouton from its receptive field location as follows: In the monocular zone, it is the orientation orthogonal to the line connecting the receptive field center and the center of vision. If the receptive field center was in the binocular zone, the preferred orientation was modeled horizontal. The coordinates of the border between the binocular and monocular zones were taken from ^50^. For every field of view, the mean retinotopic location and preferred orientation of all boutons was compared to the corresponding preferred orientation of the model. The prediction error consisted in the angular difference between the mean preferred orientation of OS boutons in a field of view, and the aforementioned modelled colliculus orientation.

### Creation of direction maps

The map of polar distributions of DS bouton’s preferred direction was computed by binning boutons based on their RF center. The size of the bins were 15° elevation and 20° azimuth. At the center of each bin, we plotted the polar histogram of DS boutons belonging to that bin. The angle bin size for the polar histograms was 15°. We overlayed four arrows which angle and length corresponded to the mean preferred direction and fraction of DS boutons, respectively, for each of the four cardinally labelled DS boutons (see classification below).

Flow fields direction maps consisted of the mean preferred direction of DS boutons labelled as either upward or posterior for every field of view (see classification below). The arrow bases coincide with the coordinates of the mean receptive field center of boutons in each the field of view. The length of the arrow indicates the percent of upward or posterior motion preferring boutons among all DS boutons in the field of view.

#### Classification of preferred direction into one of the four cardinal directions

DS boutons were classified into one of the four cardinal directions based on their preferred direction. The [-180°, 180°) space was considered, where -180° and 0° correspond to the anterior and posterior directions, respectively, and -90° and 90° correspond to the downwards and upwards direction respectively. DS boutons were classified as tuned to anterior motion if their preferred direction was included in the interval [-180°, -150°) or [120°, 180°). Temporal DS boutons were included in the [- 90°, 36°) interval. Upwards DS boutons were in [36°, 120°). Downwards in [-150°, -90). Intervals were determined manually by observing the cut-offs of the preferred direction distribution in figure 6A.

#### Quantification of CART+ ganglion cells

Confocal image stacks of the anti-CART and anti-GFP stained retinas were opened in Fiji and the number of GFP-positive and double-positive cells (CART/GFP) was counted by overlaying the two color channels. Cells were marked using the point tool and counted manually using the cell counter plugin. Significance between unilateral and bilaterally labeled CART/GFP neurons was determined by a permutation test from all three retinas per condition.

#### Quantification of preferred direction bias

Preferred directions of the recorded ganglion cells were clustered into groups by K-means clustering with periodic boundary conditions from 0° to 360°. The number of clusters for each fit were fixed by how closely the cluster centroids resembled cardinal directions (separated by 90°). A Mann-Whitney U test was used to quantify whether the number of cells in each cluster were significantly different from one another. Each cluster was fitted with a Gaussian and distributions plotted into a polar plot where height represents percent of neurons per cluster.

### Transformation of DS ganglion cells retinal coordinates into global mouse centered coordinates

To compare our data with the data acquired in the retina from ^18^, we transformed the latter data set into our global mouse centered coordinates.

#### The spherical retina coordinate system

Data from ^18^ is accessible in three dimensional cartesian coordinates on the spherical retina. Importantly, we considered here, as in ^18^, that the optical axis of the eye intercept the retina at the blind spot. Each ganglion cell body location is defined by its X, Y and Z coordinates on the retina. The (0, 0, 0) point corresponds to the blind spot. The X axis connects the nasal and temporal locations of the retina, positive values being more nasal. The Y axis coincides with the dorsal - ventral axis of the retina, positive values being more dorsal. The Z axis coincides with the optical axis of the eye, positive values being closer to the nodal point of the eye. Each ganglion cell’s preferred direction is defined by a unitary vector of coordinates U, V, and W.

#### The global mouse centered coordinate system

Our global mouse centered coordinate system was defined by 2 variables, azimuth and elevation, and 2 reference planes, the horizontal and vertical. The horizontal plane (elevation 0°) was orthogonal to the gravitational axis and intercepting the nodal eye points. The vertical plane (azimuth 0°) was parallel to the gravitational axis and intercepting the nose and the midpoint between the two eyes. Importantly, we considered the mouse’s head not in an ambulatory position, but such that the nose belonged to the horizontal plane. This puts the Lambda-Bregma axis of the skull at an approximate null angle relative to the anterior - posterior axis of the mouse. In this conditions, the optical axis of the eye lies a - 7° elevation (22° – 29°) and 64° azimuth ^51^. The azimuth of a point in the visual field corresponded to the angle between the vertical plane and a parallel plane intercepting this point and the midpoint between the two eyes. The elevation of a point in the visual field corresponded to the angle between the horizontal plane and a parallel plane intercepting the point and the midpoint between the two eyes.

#### Conversion from spherical retina coordinates to global mouse centered coordinates

Given the two above descriptions, to transform ganglion cell locations in spherical retinal coordinates into predicted receptive field center locations in our global mouse centered coordinates, we applied the following two formulas:

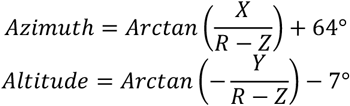

The radius R corresponded to the inner radius of the retina defined by the spherical geometry of ganglion cell body locations. We approximated this value from the data of ^18^, by averaging half the distances between the pairs of ganglion cells the farthest apart on the retina. Thereby, we approximated R to 0.95 mm.

To transform the preferred directions of ganglion cells from spherical retina coordinates into our global mouse centered coordinates, we considered the points on the retina resulting from the displacement of the ganglion cell locations when applying the corresponding preferred direction vector. We then transformed these displaced points into predicted receptive field center locations using the formulas above. Finally, we computed the normalized displacement vectors of predicted receptive field centers from each pair of ganglion cell and corresponding shifted locations, and thereby obtained the vector of preferred direction in azimuth and elevation coordinates.

---

**Figure S1.**
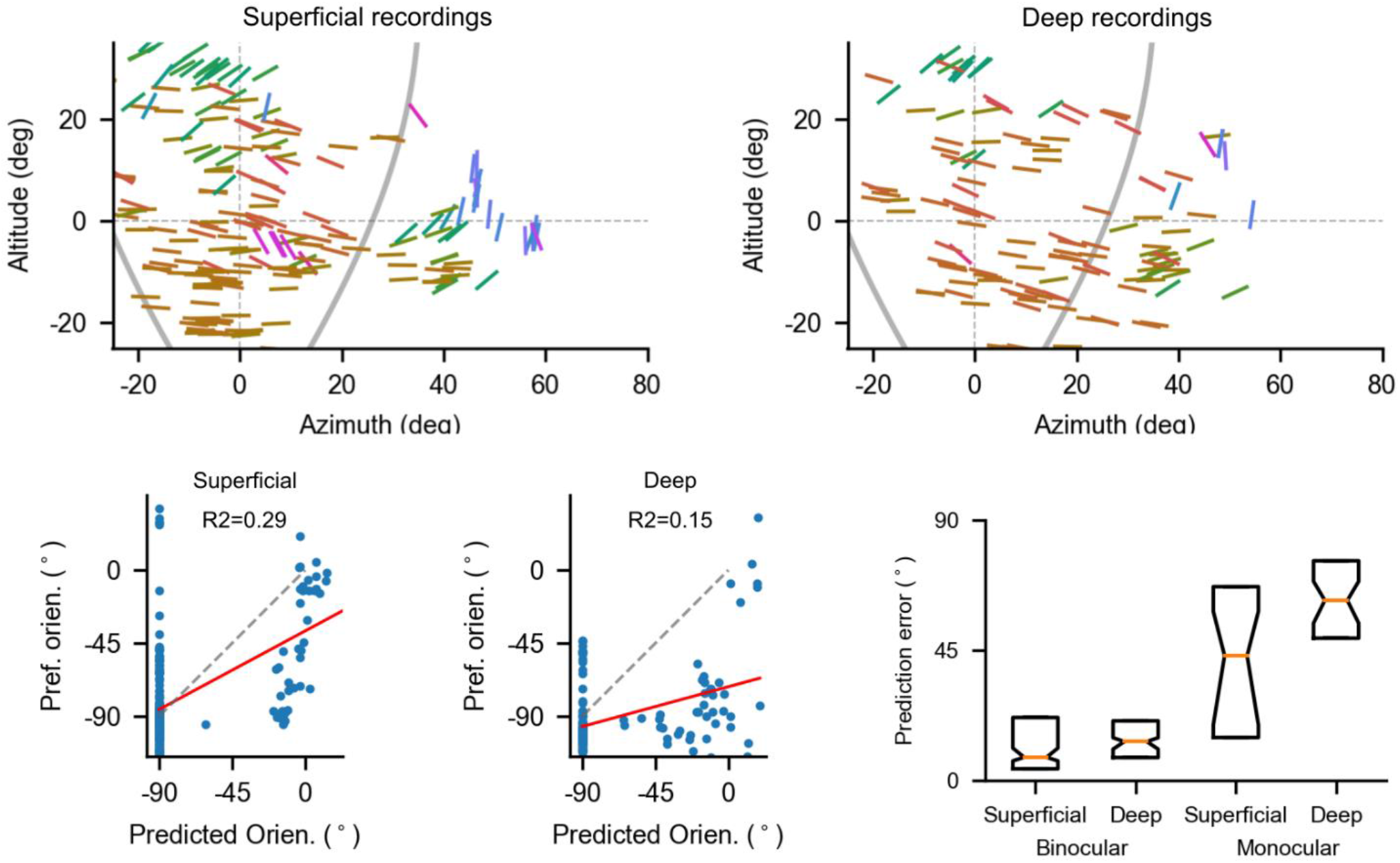
related to Figure 3: The orientation map of superficial and deep OS SC neurons. **A-B**. Retinotopic maps of preferred orientations of OS SC neurons of the most superficial (A, depth ≤ 60 μm) and deep layers (B, depth > 60 μm). **C-D**. Model comparison. Preferred orientation of each patch in superficial (C) and deep layers (D) is compared to its concentric model prediction obtained from receptive field location (see Methods). Model: grey dashed line; linear fit: red line. **E**. Prediction error between preferred orientation and concentric prediction was calculated for each cell. Boxplots show median and interquartile range of superficial and deep layer neurons, divided further into binocular and monocular zone. Notches indicate confidence interval of the median from bootstrapping. Data from (de Malmazet et al., 2018).

**Figure S2.**
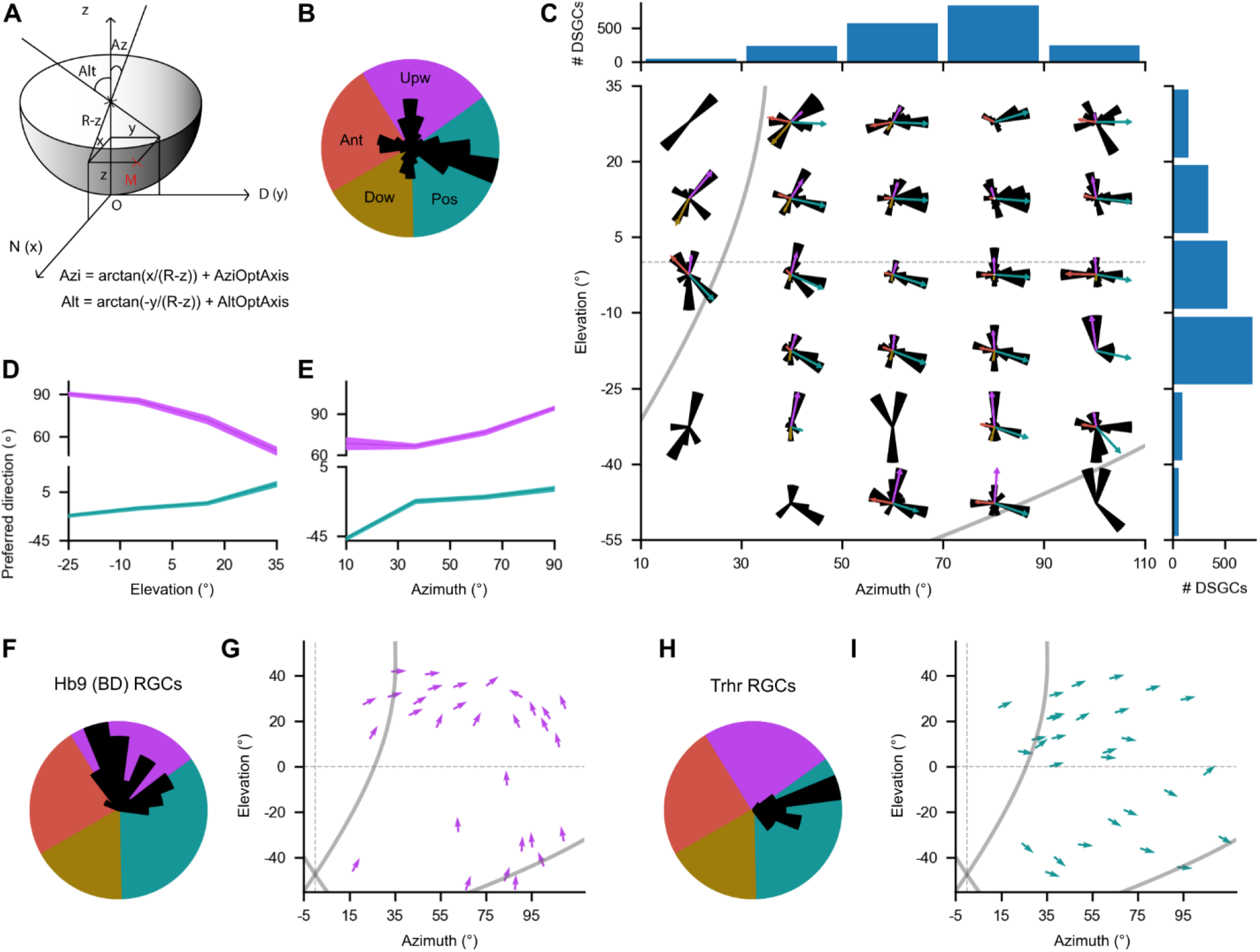
related to Figure 5:Retinotopic map of preferred direction for DS ganglion cells in global mouse-centered coordinates (Data from Sabbah et al., 2017, Ref 18) **A**. Schematic of the transformation of retinal coordinates into visual field coordinates (See methods for details). **B** Polar distribution of preferred directions for all ON-OFF DS ganglion cells (n=1953). Color wheel indicates range of each cardinal direction: anterior (red), upward (purple), posterior (teal) and downward (yellow). **C** Polar distributions of preferred directions as in (B) at equidistant receptive field locations. Arrows indicate weight and mean preferred direction of each cardinal direction. Blue histograms indicate the number of DS ganglion cells at each location. **D-E:** Mean preferred direction of upward (purple) and posterior motion (teal) preferring ON-OFF DS ganglion cells by elevation (D) and azimuth (E). **F-I:** Polar histograms (F and H) and flow fields (G and I) of preferred directions for two genetically defined ON-OFF-DS subtypes: Hb9 ganglion cells (F and G, n=31) and Trhr ganglion cells (H and I, n=31). Arrow angles and base coordinates reflect the preferred direction and retinotopic location of each cell, respectively.

